# Glial expression of *Drosophila UBE3A* causes spontaneous seizures modulated by 5-HT signaling

**DOI:** 10.1101/2024.02.08.579543

**Authors:** Megan Sleep, Saul Landaverde, Andrew Lacoste, Selene Tan, Reid Schuback, Lawrence T. Reiter, Atulya Iyengar

## Abstract

Misexpression of the E3 ubiquitin ligase *UBE3A*is thought to contribute to a range of neurological disorders. In the context of Dup15q syndrome, excess genomic copies of *UBE3A* is thought to contribute to the autism, muscle tone and spontaneous seizures characteristic of the disorder. In a *Drosophila* model of Dup 15q syndrome, it was recently shown glial-driven expression of the *UBE3A* ortholog *dube3a* led to a “bang-sensitive” phenotype, where mechanical shock triggers convulsions, suggesting glial *dube3a* expression contributes to hyperexcitability in flies. Here we directly compare the consequences of glial- and neuronal-driven *dube3a* expression on motor coordination and neuronal excitability in Drosophila. We utilized IowaFLI tracker and developed a hidden Markov Model to classify seizure-related immobilization. Both glial and neuronal driven *dube3a* expression led to clear motor phenotypes. However, only glial-driven *dube3a* expression displayed spontaneous immobilization events, that were exacerbated at high-temperature (38 °C). Using a tethered fly preparation we monitored flight muscle activity, we found glial-driven *dube3a* flies display spontaneous spike discharges which were bilaterally synchronized indicative of seizure activity. Neither control flies, nor neuronal-*dube3a* overexpressing flies display such firing patterns. Prior drug screen indicated bang-sensitivity in glial-driven *dube3a* expressing flies could be suppressed by certain 5-HT modulators. Consistent with this report, we found glial-driven *dube3a* flies fed the serotonin reuptake inhibitor vortioxetine and the 5HT_2A_ antagonist ketanserin displayed reduced immobilization and spike bursting. Together these findings highlight the potential for glial pathophysiology to drive Dup15q syndrome-related seizure activity.

## 1. Introduction

Altered expression of the E3 ubiquitin ligase *UBE3A* in the nervous system is associated with a variety of neurological disorders (J. LaSalle et al., 2015; Lopez et al., 2019). In humans, *UBE3A* is located on 15q11.2-q13 and is paternally imprinted (Lalande, 1996; Vu & Hoffman, 1997). Maternal allele deletions and loss-of-function mutations in *UBE3A* cause Angelman syndrome (Kishino et al., 1997; Matsuura et al., 1997), characterized by cognitive impairment, developmental delay and a consistently happy demeanor (Angelman, 1965; Williams, 2005). In contrast, maternal duplications of the 15q11.2-q13 region (Dup15q syndrome), are associated with autism spectrum disorder (ASD) as well as intellectual disability, muscle hypotonia and pharmacoresistant epilepsy (Battaglia, 2008; DiStefano et al., 2016; Nora Urraca et al., 2013). Indeed, as many as 3-5% of all ASD cases are estimated to arise due to Dup15q syndrome (Depienne et al., 2009), and difficult to control seizures are a major concern for the families of patients with the disorder (Conant et al., 2014). Although many genes are located in the 15q11.2-q13 region, *UBE3A* is the only gene that is paternally imprinted (maternally expressed in neurons). Paternally derived duplications of 15q11.2-q13 may be associated with anxiety and sleep problems or show no phenotypes at all, but are rarely associated with epilepsy (Cook et al., 1997; J. M. LaSalle et al., 2015; N. Urraca et al., 2013). Given the promiscuity of UBE3A, and E3 ubiquitin ligase, in marking proteins for proteolytic degradation, the molecular pathways linking *UBE3A* overexpression to Dup 15q-associated phenotypes remain to be fully elucidated (reviewed in (J. M. LaSalle et al., 2015)).

To determine how *UBE3A* overexpression causes Dup15q syndrome, several mouse models have been created (Copping et al., 2017; Smith et al., 2011; Takumi, 2011). Mice carrying a duplication of a 6.3 Mb region syntenic to 15q11.2-q13 in humans display abnormal social interaction and behavioral inflexibility, but only when paternally inherited (Nakatani et al., 2009). Neuron-specific overexpression of *UBE3A* in mice is linked with increased anxiety-like behaviors, learning and memory deficits and hypersensitivity to chemoconvulsants (Copping et al., 2017). Although at least one model showed behavior differences after strong chemical seizure induction in mice expressing elevated *Ube3a* (Krishnan et al., 2017), none of the models thus far recapitulate the spontaneous seizure phenotypes characteristic of Dup15q syndrome. Studies in *Drosophila* indicate that overexpression of *dube3a* (the ortholog of *UBE3A*) in glia, not neurons, results in a “bang-sensitive” phenotype. The bang-sensitive seizure assay (BSA) employs a mechanical shock to trigger stereotypic patterns of spasm and paralysis indicative of seizure activity in flies (K. A. Hope et al., 2017; B. Roy et al., 2020). This seizure-like activity is reminiscent of other fly mutants (*FAK, ATPa, zydeco*,) where glial disfunction is thought to contributed to behavioral phenotypes (Hope et al., 2020). Although the fly glial expression model of Dup15q syndrome recapitulates the seizure phenotypes using the BSA, a more unbiased and automated system is needed for larger anti-epileptic drug screening using this valuable disease model system.

Here we employed the IowaFLI Tracker system to quantify movement of flies overexpressing *dube3a* in glial cells after heat induction of epileptic activity. We also investigated spontaneous seizure activity in animals expressing Dube3a in glia versus neurons using head fixed electrophysiology for the first time. Finally, we show that drugs previously shown to suppress epileptic activity in this model could also effectively suppress the spontaneous epileptic behavior.

## 2. Materials and Methods

### 2.1 *Drosophila* stocks and husbandry

The UAS *dube3a* transgenic construct (BL 90373) pan-glial *repo-*Gal4 driver (BL 7415) and pan-neuronal *neurosynaptbrevin nSyb-Gal4* driver (BL 51941) have been previously described (Faiza Ferdousy et al., 2011; Kevin A. Hope et al., 2017). All flies were reared on standard cornmeal media (Bloomington Stock Center Recipe) and kept at 25 °C (70% relative humidity) on a 12:12 light-dark cycle.

### 2.2 Lifespan and behavioral analysis

Lifespans were performed as described in (Iyengar et al., 2022). Flies were collected in a 1-day age-range under CO_2_ anesthesia and kept in standard vials. The number of survivors was counted daily, and flies were transferred to fresh vials three times per week.

Automated open-field behavior monitor was performed as described in (Iyengar et al., 2012; Ueda et al., 2023), with modifications for heating the arena (Figure 1A). Polyacrylate behavioral arenas were placed on a piece of Whatman #1 filter paper, which was in-turn placed on the Peltier temperature-controlled stage (AHP-1200CPV TECA Corporation, Chicago, IL). Arena temperature was monitored by a T-type thermocouple (5SRTC-GG-T-30-36, Omega Engineering, Norwalk, CT) connected to a data acquisition card (NI TC-01, National Instruments, Austin TX). Stage temperature was controlled by a custom-written LabVIEW script (National Instruments). A custom-built PVC lighting cylinder with an LED strip was placed above the behavioral arenas (see (Ueda et al., 2023) for details). Light intensity (∼ 1000 lx) was set by a current controller driving LEDs (∼100 mA).

**Figure 1.**
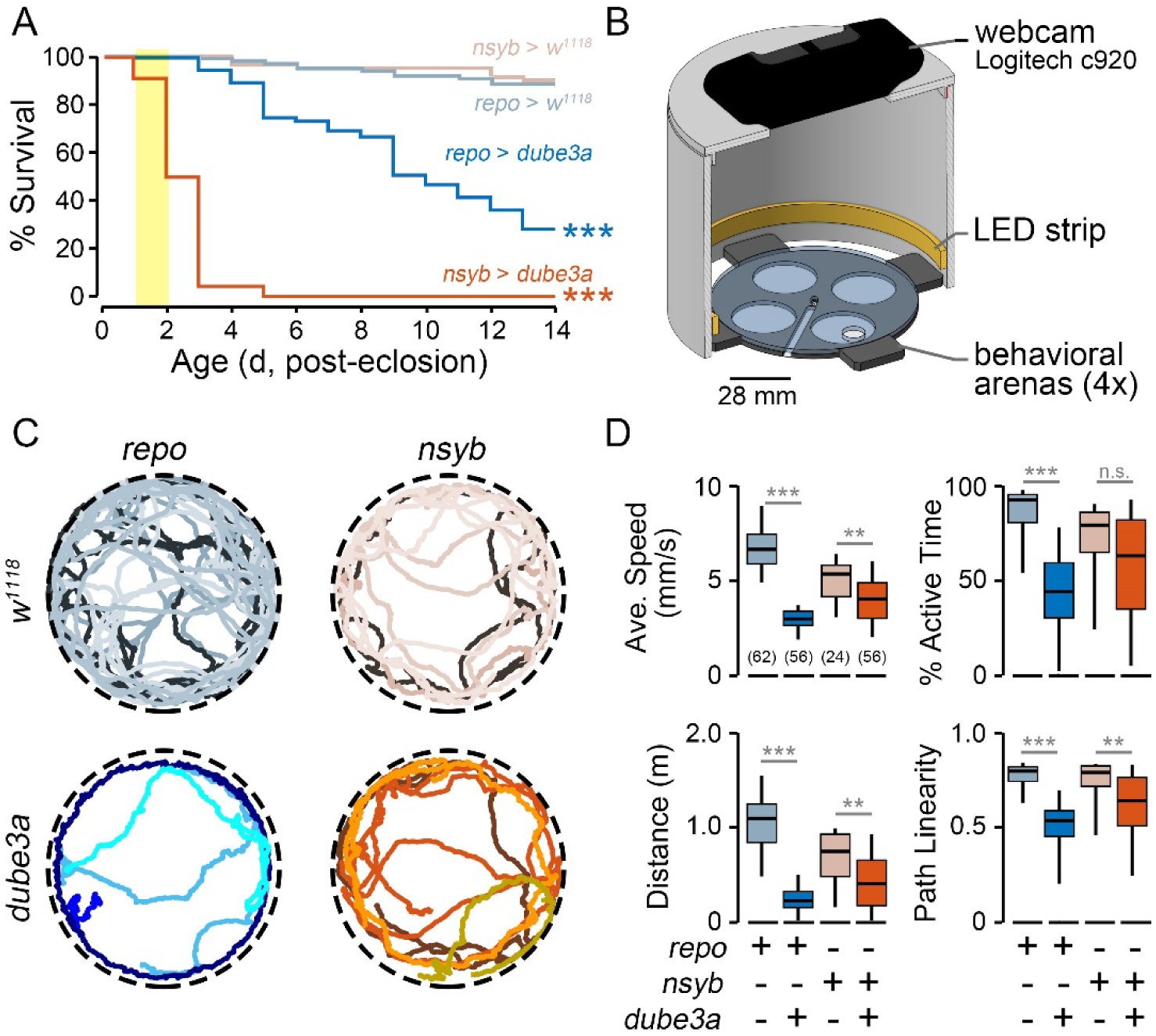
Survival and motor coordination phenotypes produced by *dube3a* overexpression. (A) Post-eclosion survival of *repo>dube3a*, *repo>w^1118^, nsyb>dube3a* and *nsyb>w^1118^* flies. 2-3 d flies were used for subsequent behavioral experiments (yellow bar). (B) Diagram of automated video tracking chamber. Up to four flies were loaded in each of the behavioral arenas. (C) Representative tracks (30-s duration) of four flies from the respective genotypes (temperature: 2°C). (D) Distributions of locomotion characteristics over the 120-s observation period for the respective genotypes. Box plots indicate the 25^th^, 50^th^, and 75^th^ %-tiles; whiskers indicate the 5^th^ and 95^th^ %-tiles. Sample sizes indicated in parentheses in the average speed panel. For lifespan analysis *dube3a* overexpression populations were compared against respective *w^1118^* controls (log-rank test). For locomotion analysis, a Kruskal Wallis non-parametric ANOVA (Bonferroni-corrected rank-sum *post hoc* test) was employed. * p ≤ 0.05; ** p ≤ 0.01; *** p ≤ 0.001.

For open-field behavior experiments, four flies of a selected genotype and sex were placed without anesthesia into behavior arenas. We employed a standardized 10-min protocol (Figure 2A) to monitor fly activity: 180 s at baseline temperature (21 °C), temperature-ramp to 36 °C (90 s), high-temperature period (36 °C-39 °C, 120 s), return to baseline (210 s). Selected 2-min periods during the baseline and high-temperature phases (30 – 150 s and 270 – 390 s) were chosen for subsequent analysis. Video recordings were made with a Logitech c920 webcam (frame rate: 30 fps) controlled by a LabVIEW script. Fly locomotion videos were analyzed using IowaFLI Tracker (Iyengar et al., 2012) running on MATLAB (R2022a, Mathworks, Natick, MA).

**Figure 2.**
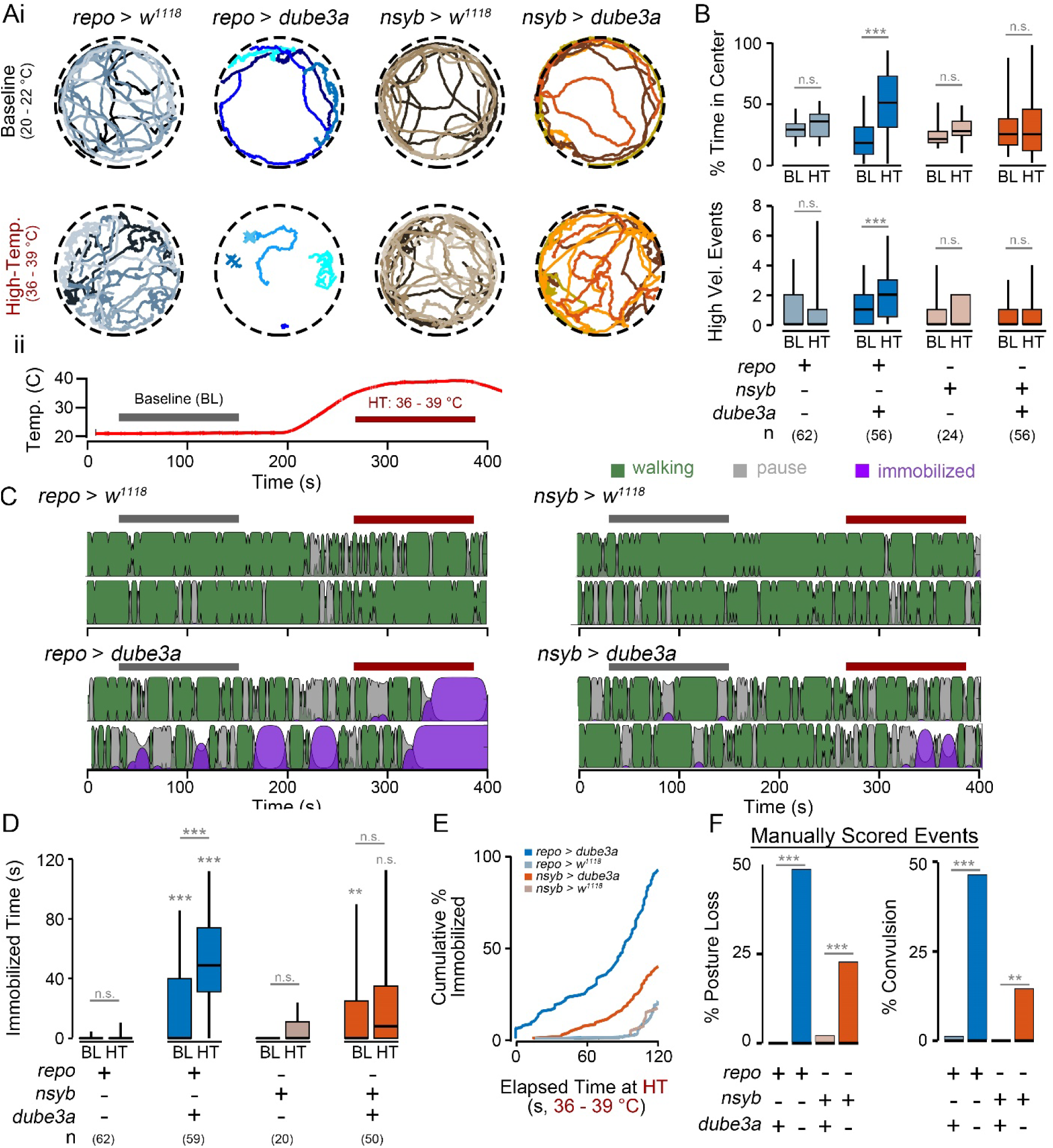
Seizure-associated motor phenotypes induced by high temperature shift in glial-and neuronal-driven *dube3a* overexpression. (Ai) Representative tracks (30-s) from *repo>dube3a*, *nsyb>dube3a* and respective controls during baseline activity (20 – 22 °C) and activity at high-temperature (36 – 39 °C). Note the abrupt and uncoordinated activity in *repo>dube3a* flies. (ii) Temperature profile corresponding to the tracks in Ai. (B) Quantification of time in center (upper panel) and high velocity events (lower panel) during the baseline and high-temperature periods. Note the increased values at high-temperature in *repo>dube3a* flies. (C) Representative classification of activity by the HMC of respective genotypes (activity from two flies is classified for each genotype). Colors indicate the classified state (walking-green, paused-pink, immobilized-purple), and the height represents the confidence of classification. Red bars indicate high-temperature period. (D) Distributions of total immobilization time during the baseline and high-temperature periods. (E) Cumulative percentage of flies classified as immobilized for > 1 s during the high-temperature period. (F) Manual scoring of posture loss and convulsions during the high-temperature period. For panels, B, and D significance determined by Kruskal Wallis non-parametric ANOVA (Bonferroni-corrected *post hoc*). For panel E, log-rank test. For panel F, Fisher’s exact test. (* p ≤ 0.05; ** p ≤ 0.01; *** p ≤ 0.001).

IowaFLI Tracker captures several locomotion parameters including average speed, percent active time, and total distance travelled as described in (Iyengar et al., 2012). The path linearity measure is described in (Chi et al., 2022), while the percent time in center is described in (Kasuya et al., 2019). High velocity events (Figure 2) were computed by finding the number times a fly’s velocity exceeded 10 SD greater than the average speed of that fly in the recording. All computations were done in MATLAB (R2023b).

Identification of “posture loss” and “convulsions” (Figure 2F) required manual scoring. Posture loss was operationally defined as a fly on its back (supine) or side or otherwise not standing on its legs. Convulsions were operationally defined as a fly displaying a wing buzz or otherwise abruptly moving in the arena while not walking. Behaviors were scored offline by a trained observer blinded with respect to the genotype.

### 2.3 Behavioral classifier construction

The behavioral classifier is a hidden Markov model (HMM) that identified periods of walking, pausing and immobilization in the activity pattern of a fly. The analysis consists of three stages: 1) pre-processing, where the full x-y trajectory of the fly is spit into sequential short (2 s) trajectories which are then aligned and scaled according to principal components; 2) classifier construction, where we created a HMM based on a training data set to determine one of three states, ‘walking’, ‘pausing,’ and ‘immobilization’, was most likely for a particular 2 s trajectory; and 3) decoding, where the HMM is used to classify trajectories from the full data sets into one of the three categories.

#### 2.3.1 Pre-Processing

To prepare tracks for classifier analysis we first segmented tracks into 2-s coordinate trajectories consisting of 61 (**x**, **y**) coordinate pairs, i.e. **x** = x_0,_…,x_60_ and **y** = y_0,_…,y_60_. Successive 2-s trajectories overlapped with the previous trajectory by 1 s. Thus, for a 10-min video containing ∼ 18,000 frames, there are ∼600 trajectories per fly. To align trajectories, the initial values were subtracted from **x** and **y** such each trajectory started at the origin, i.e. (**x’**, **y’**) = (**x**-x_0_, **y**-y_0_). These trajectories are then rotated such that they end in the same direction. The rotation angle is θ = atan (x_60_’ / y_60_’), and the aligned coordinates (**x”**, **y”**) = (**x’** cos (θ) + **y’** sin (θ), **-x’** sin (θ) + **y’** cos (θ))

The aligned coordinate trajectories, (**x”**, **y”**) are used for all subsequent analyses (S Figure XA-C). Thus for each of the ∼600 trajectories corresponding to a single fly’s activity during the 10-min video, we have a single 61 x 2 matrix (**x”**, **y”**).

To decompose trajectories into principal components, we created a matrix, **CT**, representing all trajectories from *repo* > *w^1118^*(n = 138 flies, 74,520 trajectories) and *repo > dube3a* flies studied (n = 115 flies, 62,100 trajectories). Both males and females were used in this analysis. We created this data set by transforming each of the *i =* 136,620 coordinate trajectory into a 1 x 122 array, **ct**_i_ **=** (**x”^T^**, **y”^T^**). Each row of **CT** is coordinate trajectory **ct**_i_. Thus, the coordinate trajectories from a single fly are represented by ∼ 600 rows in **CT.** In total **CT** has 136,620 total rows, 62,100 rows from *repo > dube3a* flies, and 74,520 from *repo > w^1118^*flies. We then performed principal component analysis on matrix **CT**, using the MATLAB function ‘pca’. The first four principal components (eigenvectors) corresponded with modes of locomotion (forward movement, turning, forward jitter and turning jitter), with weights (eigenvalues) corresponding with relative variation in the data set associated. Because most variation was captured by the first principal component (78%), we used this component in constructing HMM classifier. Thus the 136,620 x 122 matrix **CT** can be reduced to a forward movement vector, **fm,** a 136,620 x 1 array, with each value representing forward movement for a single trajectory.

#### 2.3.2 HMM Classifier Construction

We created a HMM classifier that determined whether a fly was in one of three “hidden” states, <u>walking</u>, <u>pausing</u>, <u>immobilized</u> based on the “directly” observed parameter of forward movement (**fm**). We found a histogram of **fm** had two modes corresponding to <u>forward</u> <u>movement</u> or <u>no forward movement</u> (with a cutoff at -11 units). To find the state transition and emission matrices of the HMM, we used the Baum-Welch algorithm, implemented in MATLAB using the ‘hmmtrain’ function. The initial transition matrix was estimated as follows:

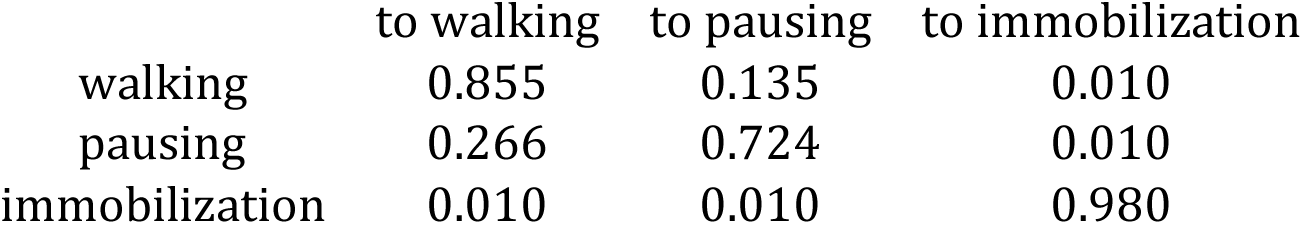

These values were based on the proportion of time *repo>w^1118^*flies displayed forward movement (65%) versus no forward movement (35%), with a low (1%) probability of transitioning to immobilization. The initial emission matrix was:

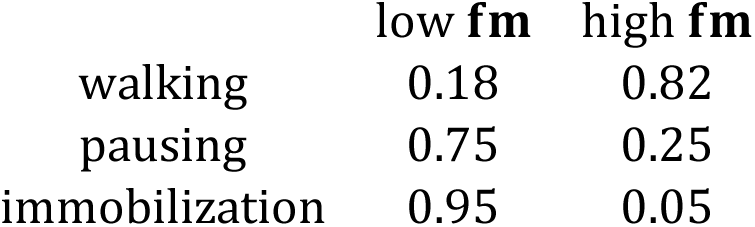

With these initial parameters, the Baum-Welch algorithm reliably converged on optimum solution. The optimal transition matrix was:

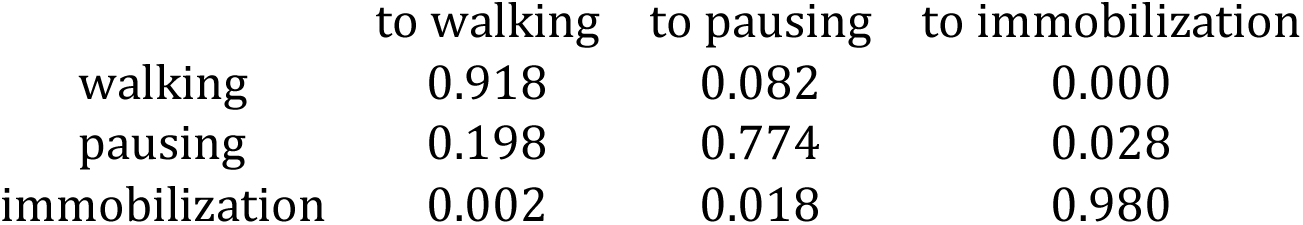

While the optimal emission matrix was:

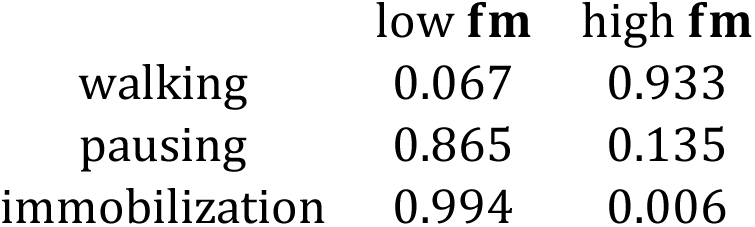

We found that varying the initial values of the emission or transition matrix by up to 20% did not alter the convergence to these optimal values.

#### 2.3.3 HMM decoding

To classify trajectories as likely walking, pausing or immobilization based on trajectories, we applied the MATLAB ‘hmmdecode’ function. The inputs to this function are the observed **fm** array, and the Transition and Emission matrices obtained following the Baum-Welch algorithm above.

### 2.4 Tethered fly electrophysiology

Electrophysiological analysis of seizure activity was based on methods described in Lee & Wu, 2002; Iyengar et al., 2014; Lee et al., 2019. Flies were anesthetized on ice and fixed to a tungsten tether pin with UV-cured cyanoacrylate glue (Loctite #4311). Following a ∼ 15 min recovery period, sharpened tungsten electrodes were inserted into the top-most left and right DLM flight muscles (DLMa) with a similarly constructed reference electrode inserted into the dorsal abdomen. Muscle action potentials were amplified by an AC amplifier (AM Systems Model 1800) and digitized by a DAQ card (NI USB 6210) controlled by a custom-written LabVIEW script. Spike trains were analyzed off-line by previously described approaches (Lee et al., 2019) implemented in custom-written MATLAB (r2023b) scripts.

Identification of burst discharges was performed as described in (Lee et al., 2019). The instantaneous firing frequency (ISI^-1^) for each spike was defined as the reciprocal of the inter-spike interval between the current spike and succeeding spike. The instantaneous coefficient of variation, CV_2_, for a pair of ISI^-1^ values *i* and *i+1* was 2*|ISI^-1^*_i_*-ISI^-1^*_i_*_+1_|/(ISI^-1^*_i_*+ISI^-1^*_i_*_+1_). Smaller CV_2_ values indicate rhythmic firing spike trains, while higher CV_2_ values indicate irregular firing. In plots of the instantaneous firing frequency versus CV_2_, bursts are readily observed as loops in the trajectory switching from burst firing (low CV_2_) to inter-burst firing (high CV_2_). A custom-written MATLAB script counted the number of loops in these trajectories to report the number of bursts.

### 2.5 Pharmacology

To block nicotinic acetylcholine receptor (nAChR)-mediated neurotransmission (Figure 3H), we used the nAChR antagonist mecamylamine (Sigma M9020). Mecamylamine was injected into the dorsal vessel (analogous to the mammalian heart) using previously described methods in the tethered fly preparation (Lee et al., 2019).

**Figure 3.**
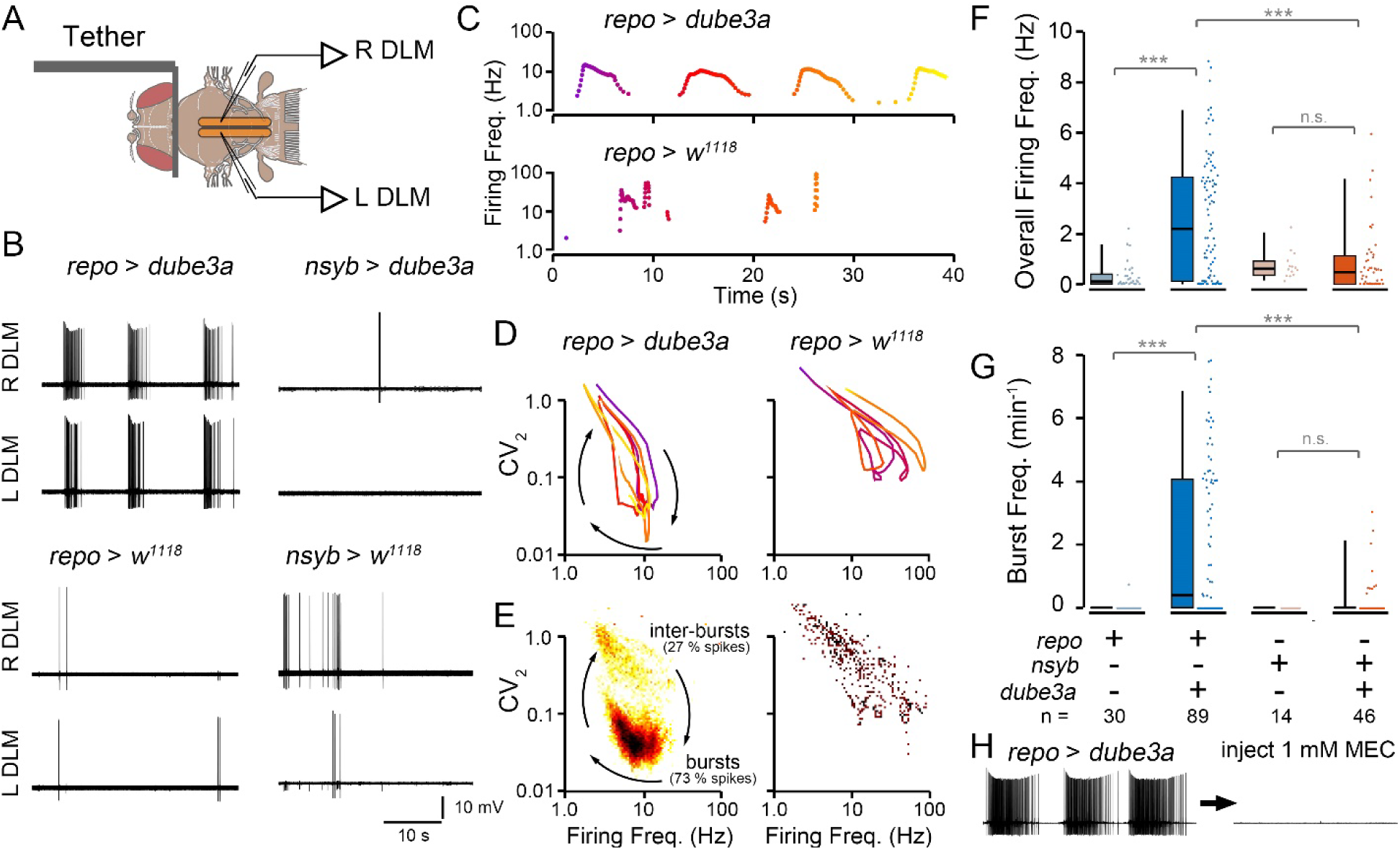
Electrophysiological monitoring of seizure activity in flies overexpressing *dube3a* in glia or neurons. (A) Illustration of the tethered fly preparation. Sharpened tungsten electrodes inserted into the left and right flight muscles (L DLM and R DLM) pick up spikes (see also S. Movie 3). (B) Representative traces of DLM flight muscle spiking from *repo>dube3a, repo>w^1118^*, *nsyb>dube3a* and *nsyb>w^1118^* flies. Note the regular bursts discharges in *repo>dube3a* that are synchronized between the left and right muscle fibers. Spiking in the other genotypes is associated with grooming activity. (C) Representative pots of the instantaneous firing frequency of spiking in *repo>dube3a* and *repo>w^1118^* flies. Color indicates elapsed time. (D) Distributions of the overall firing frequency. Sample sizes indicated above box and whisker plots. (E) Plots of the instantaneous firing rate versus instantaneous coefficient of variation (CV_2_) for the spike trains in (C). Lower CV_2_ values indicate rhythmic spiking, while higher CV_2_ values correspond with irregular firing. Note the oscillatory trajectory in *repo>dube3a*, with each loop corresponding to a single burst. (F) 2D histogram of firing trajectories in *repo>dube3a* and *repo>w1118* flies. Color indicates the number of spikes at a particular (firing frequency vs. CV2) value. Note the large number of spikes corresponding with bursts in *repo>dube3a*. (G) Quantification of the burst frequency in the respective genotypes. Bursts were not observed in the control genotypes, and only occasionally in the *nsyb>dube3a* flies. (H) Representative trace of a *repo>dube3a* fly before (left) and after (right) dorsal vessel injection of the nAChR blocker mecamylamine. Note the complete cessation of bursting activity.

The 5HT_1A_ and SERT agonist vortioxetine (VTX, HY-15414) and the 5HT_2A_ antagonist ketanserin (KET, HY-10562) were obtained from medchemexpress.com. For drug feeding experiments, VTX and/or KET were first dissolved in a stock solution (40 mM) marked with 2.5 % (w/vol) blue #1 dye. To make drug-laced media, stock solution was diluted to final concentration (0.4 µM or 0.04 µM) into melted fly food mix. Drug-fed flies were reared on VTX-or KET-laced media from larval hatching.

### 2.6 Statistical analysis

All statistical analysis was done in MATLAB using the Statistics Toolkit. A log-rank test was used to compare lifespan curves (Figure 1A), and Fisher’s exact test was used to compare relative fraction of flies displaying posture loss or convulsions (Figure 2F). All other statistical comparisons were done using non-parametric Kruskal-Wallis ANOVA following by a rank-sum *post hoc* test. P-values were corrected using the Holm-Bonferroni method.

## 3. Results

### 3.1 Distinctions in survival of glial vs. neuronal overexpression of dube3a

Previous studies in Drosophila indicated that glial, but not neuronal, overexpression of *dube3a* resulted in strong BSA phenotypes (K. A. Hope et al., 2017). Disfunction in both neuronal and glial physiology have been implicated in Dup15q syndrome, so distinguishing the specific etiology of glial vs neuronal origin is critical to the development of an effective model of the disease. To directly compare the consequences of *dube3a* overexpression in neurons versus glia, we used the Gal4-UAS system to drive expression of UAS-*dube3a* under the control of the pan-glial driver *repo-*Gal4 (*repo>dube3a*) or the pan-neuronal driver *neurosynaptobrevin-*Gal4 (*nsyb>dube3a*). We compared these flies with the respective drivers crossed with a *w^1118^*control strain (*repo>w^1118^*, and *nsyb>w^1118^*) which approximates the genetic background but lacks *dube3a* overexpression.

All crosses produced viable adult offspring. Survival of the progeny was observed over a 14-d. window (**Figure 1A**). Overexpression *dube3a* in either glia or neurons resulted in clear survival phenotypes. The median lifespan of *repo>dube3a* flies was 10 d, while *nsyb>dube3a* hand a median lifespan of only 2 d, significantly shorter than their glial expressing counterparts (p ≤ 0.001). Both overexpression populations had shorter lifespans compared to respective control flies (*repo>w^1118^* or *nsyb>w^1118^*). Thus, to ensure sufficient sample sizes, 2 d-old flies were used for behavior experiments, and 3-4 d-old flies for electrophysiological analysis. Lifespan analysis of glial and neuronal-driven overexpression of *dube3a* indicates that chronic over-expression of *dube3a* is pathological in both glia and neurons.

### 3.2 Glial overexpression of dube3a causes gross motor defects

To characterize behavioral correlates of *dube3a* expression in glial and neurons, walking activity was recorded for *repo>dube3a* and *nsyb>dube3a* flies in an open arena (**Figure 1B**). Activity was recorded via webcam and using IowaFLI Tracker to track positions (x-and y-coordinates) of each fly for quantitative characteristics of locomotion. Both *nsyb>dube3a* and *repo>dube3a* flies displayed detectable differences in walking compared to controls (**Figure 1C**). However, motor phenotypes were most extreme in *repo>dube3a* individuals. Over a 3-min observation period, *repo> dube3a* females displayed clear reductions in average speed (54.8 %), active time (52.3 %), total distance traveled (79.2 %) compared to *repo>w^1118^*controls. Although *nsyb*>*dube3a* females also displayed reductions in these parameters compared to *nsyb>w^1118^* controls, the effect sizes were relatively smaller (average speed: 24.3 %, active time: 20.4 %, distance traveled, 45.7 %), and in the case of active time were not statistically significant.

Interestingly, unlike *repo>w^1118^* controls which largely displayed straight-line or gently curved locomotion, *repo>dube3a* animals often displayed ‘jittery’ trajectories (**Figure 1C**). To quantify this feature, we developed a measure called *path linearity* which is the average ratio of displacement to distance traveled over 2-s time windows (**Figure 1D**). Consistent with the previously described BSA assays, the path linearity of *repo>dube3a* flies was markedly reduced compared to *repo>w^1118^* controls (median: 0.53 vs 0.79, p ≤ 0.001), while *nsyb>dube3a* flies displayed an intermediate effect compared to their controls (median: 0.64 vs 0.79, p ≤ 0.001). Together, these findings indicate glial expression of *dube3a* and, to a lesser extent, neuronal *dube3a* overexpression lead to disruptions in motor coordination.

### 3.3 Heat-induced and spontaneous seizure-related behavior in repo>dube3a

Previous studies indicated that seizures can be heat-induced in *repo* > *dube3a* flies (Kevin A. Hope et al., 2017). To induce aberrant behavior events without the need for mechanical shock (vortexing), *repo>dube3a*, *nsyb>dube3a*, and respective control flies in the open field arenas were subjected to a 2-min period of high temperature stress (**Figure 2A**, 36 – 39 °C). A Peltier stage below the open field arena controlled the temperature and could be rapidly changed. Consistent with previous reports, at high-temperature, *repo>dube3a* flies displayed striking seizure-associated behaviors: “wing buzzing” and “spinning” events, followed by a period immobilization, where flies would twitch or otherwise make small movements, but not walk (**Figure 2A**, **S. Movie 2**). In contrast, these behaviors were rarely observed in *nsyb>dube3a* flies and were absent in either the *repo>w^1118^* or *nsyb> w^1118^*controls.

To quantify seizure-associated behaviors, activity was compared at the baseline temperature versus high temperature using several established measures of hyperexcitable behaviors (**Figure 2B**). In open arenas, most flies tend to walk along the walls and spend little time in the arena center (‘thigmotaxis’, (Besson & Martin, 2005)), however certain hyperexcitable seizure-prone mutants (e.g. *Shudderer*) show increased time in the arena center (Kasuya et al., 2019). At baseline temperatures, both *repo>dube3a* and *nsyb>dube3a* flies (along with the control flies) spent relatively little time in the arena center. However, at high temperatures, *repo>dube3a* flies spent considerably more time in the center of the arena (p ≤ 0.001). Neither *nsyb>dube3a* flies nor the control flies displayed this increase. Similarly, there was an increase in the number of high-velocity events (corresponding with “wing-buzz” events) during high temperature in *repo>dube3a* flies, but not in the *nsyb>dube3a* or the control flies (p ≤ 0.001). Thus, glial overexpression of *dube3a*, but not neuronal expression, causes high-temperature dependent motor phenotypes reminiscent of seizure-like behavior.

### 3.4 Developing a machine-learning classifier to detect seizure-associated behaviors in flies

To establish an automated approach to identify specific moments of seizure-associated activity in *repo>dube3a* flies, we developed a machine-learning strategy. The behavioral presentation of high temperature-induced seizures in *repo>dube3a* is quite variable, with some flies displaying “spin” or “wing buzz” events, while others do not show clear “convulsion” events. However, in all cases, *repo>dube3a* flies display a prolonged period of immobilization, with leg-twitching and postural changes, but minimal movement (**S. Movie 2**). In contrast, control *repo>w^1118^* and *nsyb>w^1118^* flies either walk or engage in brief pauses in locomotion and quickly transition back to walking throughout the high-temperature period. We sought to distinguish and identify immobilization events from the normal patterns of walking interspersed with pauses which flies exhibit.

We developed a hidden Markov classifier (HMC) to identify ‘immobilization’ based on the fly’s positional trajectory. The HMC classifies a fly as ‘walking’ (high forward velocity), ‘paused’ (brief period of little forward movement) or ‘immobilized’ (prolonged period of little forward movement) in each video frame based on the fly’s locomotion characteristics at a particular moment. Control *repo>w^1118^* and *nsyb>w^1118^* flies were never classified as immobilized during normal locomotion at baseline temperatures. Details on the construction of the HMC can be found in the Methods section.

As shown in **Figure 2C-E**, during high-temperature stress, the HMC often classified *repo>dube3a* flies as immobilized (median immobilization time: 49 s), with nearly all animals (92%) classified as immobilized for at least 1 s during the high temperature period. In contrast, *nsyb>dube3a* flies occasionally immobilized (median: 10 s, 40 % of flies), with many flies active throughout the temperature stress. Importantly, repo*>w^1118^* and *nsyb>w^1118^* control flies rarely displayed immobilization events (16% and 19% of flies) with a median of only a few frames (< 30, 1-s) marked as ‘immobilized’.

Direct observation of fly behavior in individuals overexpressing *dube3a* largely corroborated findings from the HMC (**Figure 2F**). Scorers blinded to the genotype indicated ∼ 49% of *repo>dube3a* flies lost posture during high temperature stress (vs 0% of *repo>w^1118^* control flies). Similarly, 48% of repo*>dube3a* flies displayed a ‘convulsion’ (defined as a wing buzz, spinning or caroming behavior, **S. Movie 2**). Control *repo>w^1118^* flies rarely displayed this behavior (< 1%). Consistent with the HMC findings, *nsyb>dube3a* flies had a mild phenotype, with 22.4 % displaying posture loss (vs 1.9 % for control flies) and 14.7 % displaying convulsions (vs. 0 % for control flies). Together these observations indicate the HMC approach can reliably identify moments of immobilization associated with high-temperature stress, and the reported values are consistent with manual observation of seizure-associated behavior in *repo>dube3a* individuals.

### 3.5 Electrophysiological analysis of seizure activity in glial versus neuronal dube3a overexpressing flies

Given the clear seizure-like behavioral phenotypes in flies overexpressing *dube3a* in glia, we wanted to determine if more subtle electrophysiological activity signatures could be detected in these flies. We employed a tethered fly preparation previously utilized to characterize seizure activity (Iyengar & Wu, 2014). In an intact behaving fly, electrodes inserted in the dorsal longitudinal flight muscles (DLMs) enable prolonged recording of action potentials with minimal muscle damage (**Figure 3A, S. Movie 3**). In wild-type flies, DLMs display characteristically rhythmic firing during flight and arrhythmic firing during grooming behavior (Lee et al., 2019). In seizure-prone flies, a wide array of aberrant mutant-specific firing patterns have been reported (Pavlidis & Tanouye, 1995; Lee & Wu, 2002; Kaas et al., 2016; Chi et al., 2022). Importantly, DLM spiking during seizure events is correlated with hypersynchronized activity across the brain as revealed by LFP recordings (Iyengar & Wu, 2021).

In the tethered preparation, spontaneous DLM activity was monitored that was not related to flight or grooming in flies overexpressing *dube3a* in either glia or neurons and the relevant controls (**Figure 3B**). In control *repo>w^1118^*and *nsyb>w^1118^* occasional firing was identified which was correlated with grooming behaviors (**S Movie 3**). The instantaneous firing frequency during the sparse bouts (∼1-3 s) of irregular spiking approached 100 Hz (e.g. **Figure 3C**), largely consistent with previous reports of grooming-related firing (Lee et al., 2019). In sharp contrast, *repo>dube3a* flies often exhibited sustained bursts of rhythmic spiking (5-15 s), with instantaneous firing frequencies during these bursts reaching up to 20 Hz (**Figure 3C**). Across the 240 s recording period, the overall DLM firing frequency of *repo>dube3a* flies was significantly higher than the *repo>w^1118^* control individuals (**Figure 3D**, p ≤ 0.001). Overexpression of *dube3a* in neurons (*nsyb>dube3a*) did not lead to spontaneous burst discharges (**Figure 3B**). Instead, these flies displayed occasional grooming-related spiking, much like their *nsyb>w^1118^*control counterparts (**Figure 3B**). Indeed, no significant differences were observed in the overall firing frequency between *nsyb>dube3a* individuals and the *nsyb>w^1118^* control flies (**Figure 3D**).

To quantify characteristics of burst patterning in *repo>dube3a* flies, we constructed phase plots of the instantaneous firing frequency versus the instantaneous coefficient of variation, a measure of firing rhythmicity (CV_2_) (Lee et al., 2019). These plots readily differentiate the irregular grooming-associated spiking in *repo>w^1118^* flies with relatively high CV_2_ values from seizure-related burst discharges in *repo>dube3a* flies during which low CV_2_ values indicate rhythmic firing (**Figure 3E**). Histograms of firing frequency vs CV_2_ from *repo>dube3a* spiking indicated stereotypic firing frequencies (∼7 Hz) and CV_2_ values (∼0.04) corresponding with bursting (**Figure 3F**). Spiking trajectories in *repo>w^1118^* flies did not approach this region in the firing frequency – CV_2_ plots. Thus, we counted the number of times the spiking trajectory entered or exited the bursting region to quantify the frequency of burst spike discharges. Compared to *repo>w^1118^* counterparts, we found spike bursts in *repo>dube3a* flies occurred much more frequently (p ≤ 0.001), although there was wide range of burst frequencies (**Figure 3G**). Although bursting was occasionally observed in *nsyb>dube3a* flies (2/11 flies), across the cohort, there was no appreciable difference in the frequency of burst discharges from control *nsyb>w^1118^*flies.

A hallmark of seizures is hypersynchronization of neuronal activity. As shown in **Figure 3B**, in *repo>dube3a* flies, DLM spike bursts were synchronized between the left and right muscle fibers. This synchronization suggests spike bursts are centrally generated rather than arising through increased motor nerve or muscle excitability. To confirm the central origin of these bursts, we blocked central excitatory neurotransmission. Acetylcholine is the central excitatory neurotransmitter in flies, and the nicotinic acetylcholine receptor antagonist mecamylamine blocks central neurotransmission (Chi et al., 2022). We found injection of mecamylamine in *repo>dube3a* flies effectively suppressed bursting activity (**Figure 3H**). Together, these observations indicate centrally generated hyper-synchronous activity drives spontaneous spike discharges in *repo>dube3a* flies.

### 3.6 Modulation of 5-HT signaling attenuates seizure activity in flies overexpressing dube3a in glia

A previous screen for compounds ameliorating mechanical shock-induced seizure activity (BSA) in *repo>dube3a* flies uncovered several modulators of serotonin (5-HT) neurotransmission which significantly shorten the recovery time following mechanical shock (Bidisha Roy et al., 2020). In fact, Dube3a can regulate monoamine synthesis through transcriptional regulation of GTP cyclohydrolase I (F. Ferdousy et al., 2011) and in mouse models of UBE3A related syndromes, Ube3a can modulate 5HT levels in some regions of the brain (Farook et al., 2012). Elevation of overall 5-HT levels, 5-HT1A receptor agonists or antagonists of 5-HT2A receptors reduce recovery time in *repo*>*dube3a* flies, while 5-HT1A antagonists and 5-HT2A agonists increase recovery time of *repo>dube3a* flies (B. Roy et al., 2020). We selected two drugs from the previous screen, ketanserin and vortioxetine, to determine if either compound could attenuate spontaneous and heat-induced seizure-associated behavior in *repo>dube3a* flies. Ketanserin is a 5-HT_2A_ antagonist with a relatively strong effect accelerating recovery from mechanical shock, while vortioxetine is a selective 5-HT reuptake inhibitor (SSRI) with a milder but still significant recovery effect.

We found *repo>dube3a* flies fed either vortioxetine or ketanserin were resistant to high-temperature stress compared to controls on non-drug food. Specifically, during the high-temperature period of the video tracking protocol, *repo*>*dube3a* flies fed either drug at concentrations of 0.04 µM or 0.4 µM were immobilized for less time than control flies (**Figure 4A-B**). In several cases, the rate of high-velocity events was reduced as well (**Figure 4C**, 0.04 uM vortioxetine, 0.04 or 0.4 µM ketanserin). Furthermore, even at baseline temperature, spontaneous immobilization events in *repo>dube3a* flies were reduced in animals raised on vortioxetine or ketanserin food (**S. Figure 2**). Notably, neither drug completely reverses the hyperexcitability phenotypes, as immobilization and high velocity events occurred at a higher frequency in drug-treated *repo>dube3a* individuals compared to *repo>w^1118^* control flies. Together, these observations are consistent with findings from the previous BSA study, where SSRIs and 5HT2A antagonists attenuate seizure-associated behaviors in *repo>dube3a* flies.

**Figure 4.**
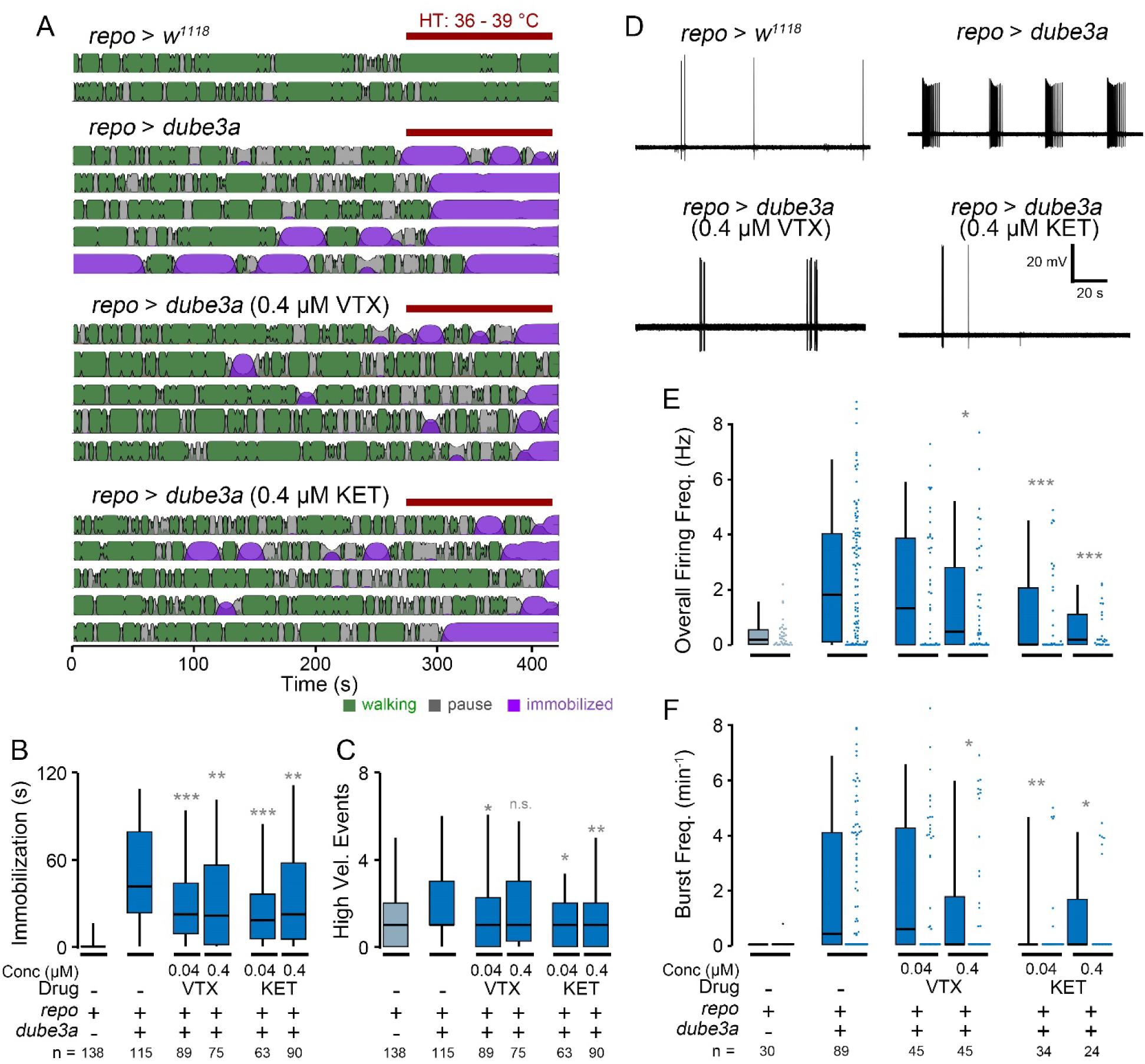
Modulation of seizure-associated behaviors and spike burst discharges in *repo>dube3a* flies with vortioxetine and ketanserin. (A) Representative activity patterns as determined by HMC in control-diet *repo>w^1118^*flies, and *repo>dube3a* flies fed either control diet, vortioxetine (VTX, 0.4 µM) or ketanserin (KET, 0.4 µM). Plots constructed as in Figure 2C, red bar indicates high-temperature period. (B-C) Box plots of (B) immobilization time and (C) high velocity events during the high-temperature period in the respective groups. (D) Representative traces of DLM spiking in *repo*>*dube3a* flies treated with VTX or KET compared to *repo*>*dube3a* and *repo>w^1118^* flies reared on the control diet. (E-F) Quantification of (E) overall firing frequency and (F) burst frequency in the respective groups. Sample seizes as indicated, statistical significance determined by Kruskal-Wallis ANOVA, rank-sum *post hoc* test. (* p ≤ 0.05; ** p ≤ 0.01; *** p ≤ 0.001).

Next, we examined whether vortioxetine or ketanserin altered the spontaneous seizure spike discharges in *repo>dube3a* flies. Consistent with the behavioral observations above, both vortioxetine or ketanserin feeding could reduce the occurrence of spontaneous burst spike discharges, and instead the spiking resembled grooming related activity observed in *repo>w^1118^*(**Figure 4D**). Specifically, vortioxetine-fed flies displayed a reduction in overall firing frequency and bursting at 0.4 µM, but not 0.04 µM (**Figure 4E & F**). Ketanserin, displayed a stronger effect, with reductions in the overall firing frequency and bursting rate detectable at both concentrations. Indeed, the overall firing frequency for Ketanserin-fed flies was not statistically distinct from *repo>w^1118^* control flies, indicating a near-complete reversal of the phenotype (p = 0.50 for 0.04 µM, p = 0.74 for 0.4 µM). These electrophysiological findings largely corroborate the behavioral findings above that SSRIs and 5HT2A antagonists can reduce seizure activity in flies overexpressing *dube3a* in glia.

## 4. Discussion

Epilepsy is a common co-morbidity in individuals with Dup15q syndrome, and seizure control is considered a major unmet medical need in these patients (Finucane et al., 1993; Conant et al., 2014). New animal models with face validity, i.e. that have spontaneous seizures, will be essential to the development of new targeted anti-epileptics in this population. Neuronal overexpression of UBE3A in mouse models of Dup15q syndrome recapitulate some aspects of the repetitive stereotypic behavior and defective social interaction characteristic of the disease (Copping et al., 2017; Takumi, 2011), however spontaneous seizure phenotypes have not been reported in these models. In flies, overexpression of *dube3a* in glia, but not neurons, causes bang-sensitive hyperexcitable phenotypes (Hope et al., 2017; Roy et al., 2020). Indeed, in the context of other epilepsy syndromes, there is growing appreciation of the role of glia in contributing to associated pathophysiology (Tian et al., 2005; Kang et al., 2005; Steinhauser et al., 2012; Patel et al., 2019). Here, we provide behavioral and new electrophysiological evidence of the critical role of glia in driving *dube3a-*associated seizure phenotypes in *Drosophila*. Importantly, this report provides the first documentation of spontaneous and recurrent seizure activity in any *UBE3A* overexpression model.

Although we found both neuronal-and glial-driven overexpression of *dube3a* led to detectable survival and motor phenotypes (**Figure 1**), *repo>dube3a* flies displayed marked seizure-associated behavioral phenotypes evoked by high temperature stress (**Figure 2**). Furthermore, electrophysiological analysis revealed most *repo>dube3a* flies showed recurrent and synchronized spike burst discharges indicative of spontaneous seizure activity (**Figure 3**). This abnormal spiking bursting was not observed in flies overexpressing *dube3a* in neurons. Lastly, we studied the effect of two previously identified compounds which attenuate bang-sensitivity in *repo>dube3a* flies (Roy et al., 2020). As shown in **Figure 4**, in *repo>dube3a* flies, we found ketanserin (a 5HT2_A_ antagonist) and vortioxetine (a SSRI) reduce both the vulnerability to high temperature stress and the occurrence of spontaneous spike discharges. Together, these findings highlight the important role of glia in driving Dup15q-related seizure phenotypes, and provide a road map for developing therapeutic strategies for the syndrome.

### 4.1 An automated approach to quantifying gliopathic seizure-related behavior in Drosophila

Like other *Drosophila* models of epilepsy, *repo>dube3a* transgenic flies are particularly amenable for high-throughput behavioral screens to identify genetic or pharmacological conditions that suppress seizure phenotypes. Two widely used methods to study seizure-associated behaviors in *Drosophila* are bang-sensitivity and high-temperature sensitivity assays, where observers score hyperexcitable behaviors following mechanical or temperature stress respectively (Benzer, 1973; Burg & Wu, 2012; Ganetzky & Wu, 1982; Melom & Littleton, 2013; Sun et al., 2012). Our approach, which combines automated fly tracking with machine learning to identify seizure-related events, makes several improvements to these established methods. First, automated fly tracking eliminates human scoring as a source of variability in the analysis. Second, the system requires fewer personnel costs, as the control of camera and stage temperature are automated. Finally, videos of fly behavior are available for *post-hoc* analysis of behavioral features well beyond the initial study.

Automated video tracking approaches have been employed to characterize walking behavior in several hyperexcitable mutants (Chi et al., 2019; Iyengar et al., 2012; Kaas et al., 2016; Stone et al., 2013). Here, we built upon this work to identify seizure-related behavioral events based on locomotion tracking using a custom machine-learning approach. Several studies have employed hidden Markov models (HMM) like the one used here to classify behavioral states in animals (e.g. (Wiggin et al., 2020); (Jiang et al., 2019)). Hidden Markov model strategies are also widely employed to detect seizure events based on EEG time-series data (Abdullah et al., 2012; Baldassano et al., 2016; Lee et al., 2018; Wong et al., 2007). We found the hidden Markov Classifier (HMC) classification was largely consistent with manual observations of immobilization behavior (**Figure 2D** & **E** vs. **Figure 2F**). Importantly the HMC provided classification on a frame-by-frame basis facilitating quantification of seizure behavior at a higher temporal resolution than the previous approach. Thus, drug-induced phenotypic variations can be quantitatively compared with each other (**Figure 4A-B**).

Despite the utility of the HMC in this study, we recognize several potential limitations in our implementation. First, the specific parameters generated for the HMC are likely specific to identifying immobilization events in *repo>dube3a* flies, as the training dataset consisted of tracks from *repo>dube3a, repo>w^1118^, nsyb>dube3a,* and *nsyb>w^1118^* flies. Future studies could optimize the model by using a larger and more diverse data set of flies exhibiting high temperature-evoked seizure-related behaviors. Indeed, there is a large number of *Drosophila* mutants which display heat induced seizures, and there are likely many subtle differences among the behavioral phenotypes displayed by these mutants. Second, our model designates ‘immobilization’ as the only seizure-related behavioral state. More sophisticated classifiers building on our approach may be used to designate the ‘spinning’ and ‘wing-buzz’ that also correspond to seizure-related behavior repertoire (**S Movie 2**). Several behavioral classification approaches utilizing high-resolution images of fly posture (e.g. (Berman et al., 2016; Mueller et al., 2019; Pereira et al., 2022)) may better capture these events for high resolution classification.

### 4.2 Electrophysiological signatures of seizure activity in dube3a overexpressing flies

Recording from DLM flight muscles of tethered flies is a convenient electrophysiological readout of seizure activity in Drosophila (Pavlidis & Tanouye, 1995; Lee & Wu, 2002; Iyengar & Wu, 2014). During seizures, Drosophila flight muscles display characteristic bursts that are synchronized between the left and right motor units (Lee et al., 2019). These bursts present as self-similar loops in plots of the instantaneous firing frequency versus the instantaneous coefficient of variation (**Figure 3D**, Lee et al., 2019). Previous studies have uncovered large-amplitude brain local field potential (LFP) signals which coincide with bursting activity (Iyengar & Wu, 2021). Our recordings revealed glial overexpression of *dube3a*, but not neuronal overexpression of *dube3a* leads to spontaneous seizures (**Figure 3B**). Approximately 50% of *repo>dube3a* flies display spontaneous seizure bursts, while only a few *nsyb>dube3a* flies and no *repo>w^1118^* or *nsyb>w^1118^*flies showed this seizure activity (**Figure 3D, E, G**). To demonstrate that the observed spike discharges in *repo>dube3a* flies originate from the central nervous system (rather than the neuromuscular junction), we injected mecamylamine, a blocker of central excitatory neurotransmission. Mecamylamine reliably blocked spike discharges indicating the activity is generated centrally (**Figure 3H**). Together, these observations suggest *repo>dube3a* flies model critical aspects of hyperexcitability observed in Dup15q patients, that of spontaneous seizure activity.

Although *repo>dube3a* represents the first case glial disfunction leading to spontaneous and recurrent seizures in flies, glial pathophysiology has been implicated in hyperexcitable phenotypes characteristic of several mutant strains. Flies carrying glia-specific disruptions of the Na^+^/K^+^ ATPase gene *ATPα* (Palladino et al., 2003), focal adhesion kinase gene *FAK* (Ueda et al., 2008) or the NCKX gene *zydeco* (Melom 2013) all display bang-sensitivity and high-temperature immobilization phenotypes like *repo>dube3a* flies. However, unlike these mutants, *repo>dube3a* flies also display spontaneous seizures at room temperature. Interestingly, the Na^+^/K^+^ ATPase pump encoded by *ATPα* was previously shown to be a potential substrate for Dube3a in flies. Furthermore, *repo>dube3a* flies exhibit high concentrations of extracellular K^+^ compared to control flies (Hope et al., 2017). Thus, a sub-set of the *repo>dube3a* seizure phenotypes may be due to attenuated *ATPα* function. However, because *ATPα* loss-of-function mutants do not show spontaneous seizures at room temperature, it is likely that other ubiquitin substrates regulated by Dube3a in flies contribute to the development of seizures. Ongoing efforts to characterize proteomic changes associated with *dube3a* overexpression may uncover additional Dube3a substrates which contribute the phenotype in *repo>dube3a* flies.

### 4.3 A pipeline to identify anti-epileptic drugs for the treatment of Dup15q syndrome

A particular strength of Drosophila epilepsy models is the high-throughput nature of behavioral phenotyping. Screens for genetic factors or pharmacological compounds that modify fly seizure phenotypes are straightforward and cost-effective (e.g. Stillwell et al., 2006). Several prior efforts have established certain anti-epileptic drugs (AEDs) can reduce susceptibility to mechanical shock or accelerate recovery in bang-sensitive mutants (Keubler & Tanouye 2002; Reynolds et al., 2003). Fly epilepsy models also offer the opportunity to identify new compounds (Kasuya et al., 2019), natural products (Dare et al., 2021) or even repurposed drugs (Roy et al., 2020) which suppress seizure activity. Interestingly, AED compounds can be efficacious in certain contexts, but have no effect on other models. For example, studies on the Na^+^ channel mutant *Shudderer* revealed that milk lipids can suppress spontaneous seizures, but these same lipids have no significant effect on the spontaneous leg-shaking and neuromuscular excitability phenotypes in the K^+^ double-mutant *eag Sh* (Kasuya et al., 2019). This observation highlights the need to develop models of a variety of pathophysiological processes leading to epilepsy and related seizure disorders.

The drug modulation studies of seizures in *repo>dube3a* flies presented here was motivated by a prior unbiased screen of FDA-approved compounds that revealed several compounds that could suppress BSA in this model (Roy et al., 2020). The Roy *et al*. screen indicated compounds affecting serotonergic or dopaminergic signaling could suppress seizure phenotypes, somewhat surprisingly since these compounds are typically used as anti-depressants, not anti-epileptics. Specifically, SSRIs, 5-HT1_A_ agonists or 5-HT2_A_ antagonists reduced the recovery time from mechanical shock in *repo>dube3a* flies significantly. In contrast, 5-HT1A antagonists or 5-HT2A agonists hindered recovery following seizure induction (Roy et al 2020). In **Figure 4**, we tested two serotonergic compounds, vortioxetine (an SSRI), and ketanserin (a 5HT2A antagonist), using the automated video tracking and tethered fly electrophysiological assays. Consistent with bang sensitivity assays in Roy et al., we found at high temperature, both drugs reduced time spent immobilized (**Figure 4B**). In the tethered fly preparation, we found both drugs (0.4 µM) could reduce both overall flight muscle spiking as well as spike bursts (**Figure 4D-F**).. Notably, most drug-fed flies eventually displayed high-temperature induced immobilization, and in these flies spontaneous spike discharges were sometimes observed. Thus, it is conceivable future drug screening efforts will yield compounds which further reverse seizure-related phenotypes in the *repo>dube3a* overexpression model of Dup15q syndrome.

Here we have shown that our gliacentric Drosophila model of Dup15q epilepsy continues to be a robust tool for the evaluation of new compounds specifically targeted to this disorder, where pharmacoresistant epilepsy is a major factor in the quality of life of these individuals. Moreover, we have now used this model to develop a robust video tracking tool to detect and quantify seizure events that we then validated using head fixed electrophysiology. The implications of this work broad given the large number of fly homologues to human epilepsy associated genes (REF). We anticipate that applying the tools developed here to other fly epilepsy models will narrow the range of drugs for specific epilepsy treatments (i.e. personalized medicine) especially using the drug repurposing approach.

## Acknowledgments

We thank members of the Iyengar and Reiter labs for their helpful discussions and technical assistance. We are grateful to Dr. Chun-Fang Wu (Univ. Iowa) for assistance during initial stages of this project, and James Pugh for constructing behavioral arenas. This work was supported by institutional funds from the University of Alabama and R01 NS115776-0A1 to LTR.

